# Prohibitin complexes associate with unique membrane microdomains in cells

**DOI:** 10.1101/2025.11.14.688579

**Authors:** Michaela Medina, Hamidreza Rahmani, Ya-Ting Chang, Benjamin A. Barad, Danielle A. Grotjahn

**Author notes:** **To whom correspondence should be addressed:** Danielle A. Grotjahn, Department of Integrative Structural and Computational Biology The Scripps Research Institute, La Jolla, CA 92037.

## Abstract

Prohibitins are multi-subunit protein complexes implicated in a variety of roles in mitochondria, including maintaining cristae architecture, regulating lipid metabolism, and mediating protein quality control. Prohibitins form large, ring-like structures within the mitochondrial inner membrane, and it has been proposed that this unique architecture enables them to organize distinct membrane environments that support their diverse functions. However, the molecular mechanisms driving these structure-function relationships within the native cellular context remain poorly defined. In this work, we combine cellular cryo-electron tomography, subtomogram averaging, and *Surface Morphometrics* analysis to systematically characterize the structure and local membrane environment of prohibitin complexes in mouse embryonic fibroblast (MEF) cells. We show that prohibitins are enriched in the inner boundary membrane subdomain of the inner mitochondrial membrane. Moreover, we find that a subset of prohibitin complexes exhibits a unique association with a matrix-facing density that resembles an *m*-AAA protease. Further, we show that prohibitins consistently localize to regions of reduced lipid bilayer thickness across all inner mitochondrial membrane subdomains, suggesting that they locally remodel the membrane to support their function. Together, these findings provide direct structural evidence that prohibitin complexes are associated with unique membrane microdomains in cells and establish a framework for uncovering how the local cellular context enables prohibitin function.

## INTRODUCTION

Mitochondria are multifaceted organelles that support a variety of eukaryotic cell processes, including energy production and metabolism, cell signaling, and stress adaptation. Their ability to perform these diverse functions fundamentally depends on their dynamic behavior, which links changes in the structure of the distinct inner and outer mitochondrial membranes (IMM and OMM, respectively) to functional adaptations in response to changing cellular demands (Tábara, Segawa et al. 2025). In particular, the IMM exhibits extensive structural complexity through the formation of cristae, the specialized subcompartments that contain the oxidative phosphorylation (OXPHOS) machinery. The presence of cristae subdivides the IMM further into distinct subdomains, including the inner boundary membrane (IBM), the cristae junction (CJ), and the crista body (CB). The formation of these structurally and functionally distinct subdomains is dictated by the unique combination of lipids and embedded proteins that sculpt or stabilize the IMM. For example, protein complexes such as the <u>mi</u>tochondrial inner <u>c</u>ontact site and <u>o</u>rganizing <u>s</u>ystem (MICOS), OPA1, and ATP synthase work in concert with mitochondria-specific lipids to remodel the IMM and promote the compartmentalization necessary for efficient ATP production (Caron and Bertolin 2024).

In addition to these well-established membrane-shaping factors, prohibitin complexes have emerged as key regulators of IMM architecture and composition (Osman, Haag et al. 2009) (Merkwirth, Dargazanli et al. 2008). Prohibitins are made up of two conserved proteins, prohibitin-1 and prohibitin-2 (PHB1 and PHB2, respectively), which are members of the SPFH (stomatin-prohibitin-flotillin-HflK/C) family that assemble into large membrane-embedded complexes and are found across all kingdoms of life. Prohibitins are indispensable for several mitochondrial functions, including mitochondrial DNA organization (Kasashima, Sumitani et al. 2008), respiration (Osman, Wilmes et al. 2007, Schleicher, Shepherd et al. 2008, Ande, Nguyen et al. 2014), biogenesis (Liu, Lin et al. 2012), stress response (Da Cruz, Parone et al. 2008), apoptosis (Fusaro, Dasgupta et al. 2003, Merkwirth, Dargazanli et al. 2008, Liu, Ren et al. 2009, Muraguchi, Kawawa et al. 2010), and mitophagy (Alula, Delgado-Deida et al. 2023). Loss of prohibitins causes severe mitochondrial defects, including mitochondrial fragmentation, aberrant cristae, and reduced respiratory capacity, suggesting that their presence helps to maintain the composition and structural integrity of the IMM. Consistent with this, previous work has shown that prohibitin complexes can regulate the stability and activity of IMM-localized OXPHOS machinery through both direct binding interactions and indirect mechanisms, including modulation of proteolysis by directly interacting with inner mitochondrial membrane proteases (Steglich, Neupert et al. 1999). Prohibitins can also influence IMM lipid composition, including the maturation of cardiolipin, a mitochondria-specific lipid essential for maintaining cristae shape and membrane stability (Lourenço, Rodríguez-Palero et al. 2021). However, the mechanisms by which prohibitin mediates its multifaceted roles within the IMM remain poorly defined.

Recent studies have leveraged advances in cellular cryo-electron tomography (cryo-ET) to directly visualize the structural architecture and localization of prohibitin complexes within native cellular environments, revealing that prohibitins form a ring-like structure within the inner mitochondrial membrane (Lange, Ratz et al. 2025, Rose, Herrmann et al. 2025, Waltz, Righetto et al. 2025). It has long been postulated that the ring-like structure of prohibitin complexes serves as a scaffold for the recruitment of specific proteins and lipids, thereby forming unique membrane microdomains within the IMM (Osman, Merkwirth et al. 2009). However, direct experimental evidence for the formation of a membrane microdomain associated with prohibitins has not been tested.

Here, we combine these latest advances in cellular cryo-ET with *Surface Morphometrics* tools and discover that prohibitin complexes are associated with unique membrane environments in mammalian cells. We show that prohibitins are localized throughout the IMM but are specifically enriched in the IBM subdomain. Interestingly, we find that a subset of prohibitins contains an extra matrix-facing density, which we propose represents a *m*-AAA protease. Across all subdomains, prohibitin complexes induce a local thinning of the lipid bilayer that is distinct from its surrounding environment. Our findings offer clear structural evidence that prohibitins establish discrete membrane microdomains through specific subdomain localization, protein associations, and targeted alterations to their local lipid bilayer environment.

## RESULTS AND DISCUSSION

### Prohibitin complexes are enriched in the inner boundary membrane (IBM) subdomain

While recent cryo-ET studies have examined prohibitin structure in *Chlamydomonas reinhardtii* (Waltz, Righetto et al. 2025) and human cancer cell lines (Lange, Ratz et al. 2025, Rose, Herrmann et al. 2025), how their structures influence the surrounding mitochondrial environment remains poorly defined. To address this, we re-analyzed our recently deposited (EMPIAR-13056) cryo-ET dataset of cryo-focused ion beam-milled mouse embryonic fibroblast (MEF) cells to define the local environment of prohibitin complexes in the native cellular context. Within the resulting tomograms, we observed the characteristic dome-like shape of prohibitins, consistent with previous studies (***Figure 1A***). To further resolve the molecular details of these complexes, we manually selected 665 individual prohibitin complexes and subjected them to subtomogram averaging (see **Methods** and ***Supplemental Figure 1, Figure 1B&C***). The resulting EM map, resolved at 34 Å with C1 symmetry, showed a hollow, dome-shaped structure protruding from the membrane, with 11 ‘spoke-like’ densities exhibiting uneven spacing (***Figure 1C, Supplemental Figure 1C***). These structural features are consistent with a recent study that reported lateral openings in the ‘walls’ of the prohibitin complex (Hong, Guan et al. 2025). Rigid body docking of the atomic model from this structure revealed strong agreement with our subtomogram structure (***Figure 1D***), suggesting that this may represent the primary structural state of prohibitins within mitochondria in MEF cells.

**Figure 1.**
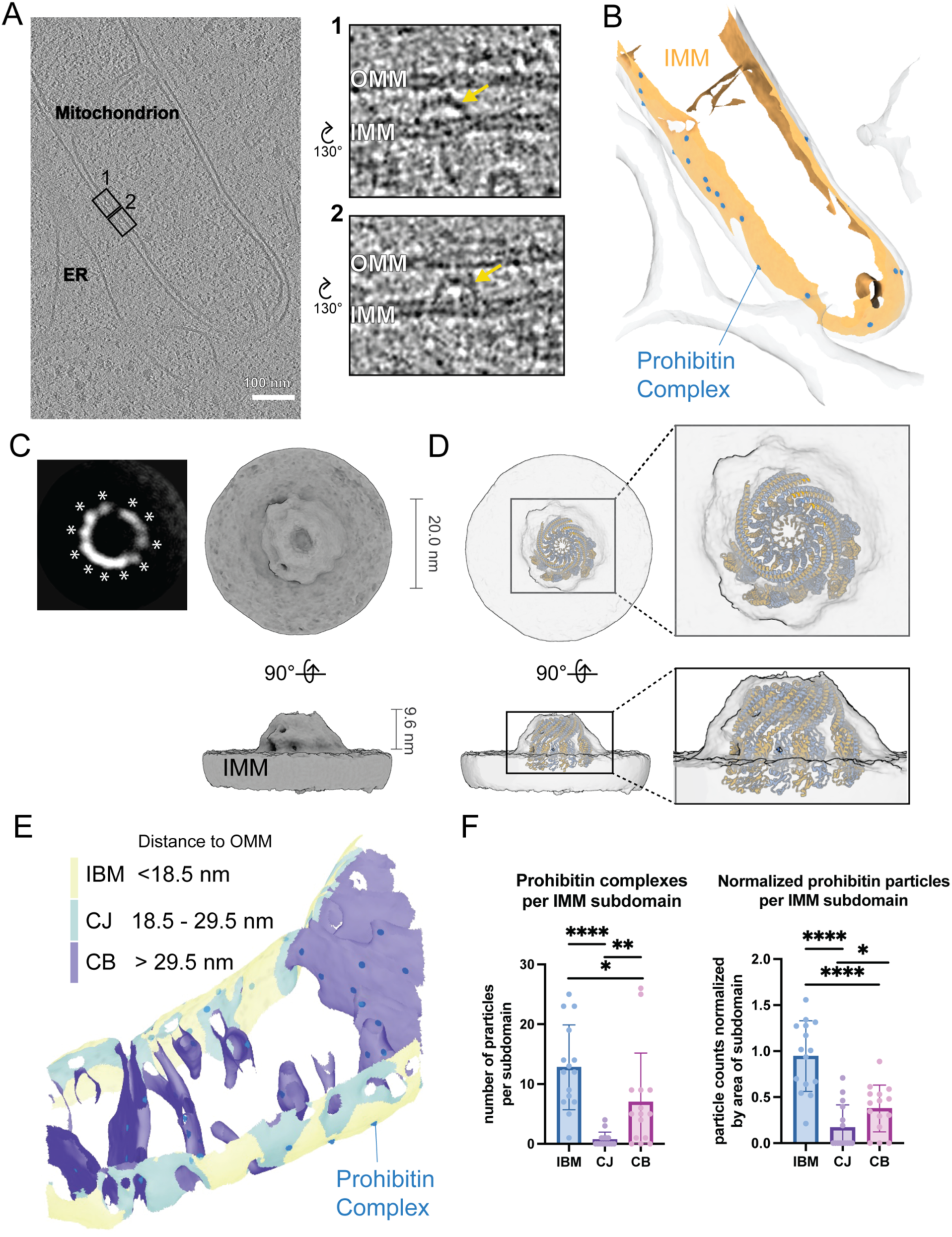
Prohibitin complexes are enriched in the inner boundary membrane (IBM) subdomain in mouse embryonic fibroblast (MEF) cells. A. Tomogram of a cryo-focused ion beam (cryo-FIB) milled lamella of a MEF cell showing a mitochondrion and surrounding endoplasmic reticulum (ER). Insets labeled 1 and 2 show individual prohibitin complexes resembling ‘dome-like’ structures (yellow arrow) that are anchored in the inner mitochondrial membrane (IMM) and extend toward the outer mitochondrial membrane (OMM). B. Triangulated surface meshes of organelle membranes visible in the tomogram in (A). The IMM is shown in orange with mapped coordinates of the prohibitin complexes overlaid in blue. The OMM and ER membranes are colored transparent gray. C. A 2D slice of the subtomogram average structure of the prohibitin complex from 665 subvolumes showing clear distinctions between prohibitin heterodimers (white asterisks, left panel). Two views of the subtomogram average of the prohibitin complex are shown as a gray isosurface rendering (right panels). D. Two views of the subtomogram average (gray transparent density) of the prohibitin complex with fitted atomic models of prohibitin in the ‘open’ conformation (PDB-9UNL). E. Triangulated surface mesh of the IMM colored by IMM-OMM distance to label the distinct subdomain: inner boundary membrane (IBM, yellow), cristae junction (CJ, teal), and cristae body (CB, purple). The IMM mesh is overlaid with the prohibitin particle positions overlaid in blue. F. Quantification of the number of prohibitin complexes within each subdomain (left) and the particles normalized by the total area of the subdomain per mitochondrion (right). n = 15 tomograms. P values from Mann-Whitney U test are indicated *P< 0.05; *P< 0.05;***P<0.001; ****P<0.0001

A major advantage of cellular cryo-ET over other structural methods is that it can capture both the native structure of macromolecules and their spatial localization within intact cellular environments. We took advantage of this capability to assess whether prohibitins exhibit preferential localization to specific IMM subdomains (e.g., IBM, CJ, CB). We used a distance-based filtering strategy within our *Surface Morphometrics* pipeline to define the distinct inner membrane subdomains based on their proximity to the outer membrane (***Figure 1E***) (Barad, Medina et al. 2023). Next, we identified the nearest triangle on the membrane surface for each refined prohibitin particle coordinate, thereby assigning each individual prohibitin complex an IMM subdomain identity. Using this structure-mapping analysis, we found that, although prohibitins are localized to some degree across all domains in the inner membrane, there was a significant increase in the number of prohibitins localized to the IBM regions (***Figure 1F***). This is in contrast to previous reports from human cancer cell lines (e.g., U2OS cells) (Lange, Ratz et al. 2025), which report that the majority of prohibitin complexes localize to the CB. These differences may reflect variations in cellular metabolic state between cancer and fibroblast cells or species-specific differences. Given that IBM is enriched in mitochondrial protein import and proteolytic machinery, enrichment of prohibitins within this subdomain suggests that prohibitins may play an active role in protein homeostasis in this cellular context.

### A matrix-facing density is associated with a subset of prohibitin complexes

Prohibitins interact directly with *m*-AAA proteases, such as AFG3L2 and paraplegin (SPG7), forming a supercomplex that regulates the turnover of unassembled respiratory chain components (Steglich, Neupert et al. 1999). Given that the majority of the prohibitin complexes in our data reside in the IBM, we wondered whether they exhibit additional structural features suggestive of an active role in mitochondrial proteolysis or protein import. To assess this, we expanded the box size of our subtomogram average and performed a masked 3D classification on the matrix-facing side of the density in the IMM region just below the IMS-facing prohibitin “dome” (***Supplemental Figure 1A&B***). This analysis yielded two structural classes, both showing apparent density for the membrane and prohibitin complex, with one class (Class 1) displaying a prominent additional density on the matrix side of the IMM and a separate class (Class 2) devoid of this additional density (***Figure 2A***). Notably, Class 1 shares striking structural similarity to the two-dimensional negative-stain class averages reported in a recent study of a purified, *in vitro* reconstituted prohibitin-*m*-AAA-protease complex (Luo, Zheng et al. 2025). Although the resolution is insufficient for unambiguous identification, fitting an atomic model of the *m*-AAA protease AFG3L2 (Puchades, Ding et al. 2019) into this region in Class 1 structure produced good overlap with the underlying density as assessed by visual inspection (***Figure 2B***), suggesting that this density may correspond to an IMM protease or another complex of equivalent size and shape. Visual inspection of individual prohibitin complexes from Class 1 showed similar, albeit less well-defined features in this region, suggesting that the signal is present in the raw data and unlikely to be an averaging artifact given the crowded nature of the mitochondrial matrix environment (***Figure 2C***). We further validated the presence of this additional matrix-facing density in the raw tomograms using patch-based density line-scan analysis, comparing membrane patches corresponding to particles within the two distinct classes (Medina, Chang et al. 2025) (***Figure 2D***). The profiles for both Class 1 and Class 2 revealed two prominent peaks corresponding to the inner membrane bilayer and an additional peak ∼15 nm away on the side of the intermembrane space (IMS) toward the outer membrane, consistent with the signal coming from the top of the prohibitin “dome”. However, only the Class 1 line-scan profile displayed an additional, prominent peak at ∼10 nm from the bilayer midline on the matrix side. These results show that a subset of prohibitin complexes in MEF cells is associated with an additional matrix-facing density positioned opposite the IMS-facing dome portion of the prohibitin complex. Based on its dimensions and localization, we propose that this density represents a *m*-AAA protease.

**Figure 2.**
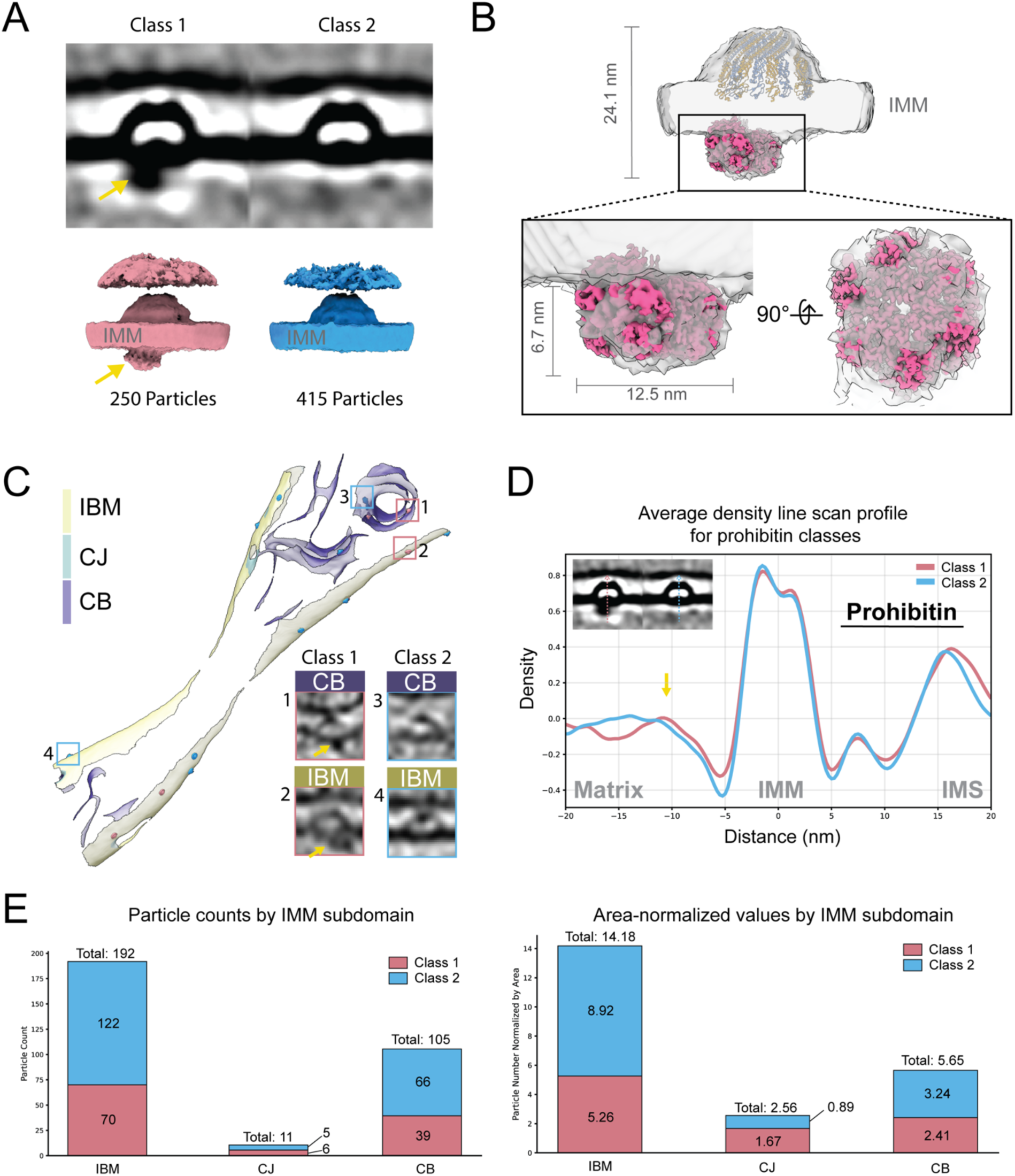
A matrix-facing density is associated with a subset of particles in the subtomogram average structure of the prohibitin complex. A. 2D slices (top) and isosurface renderings (bottom) showing two separate classes of particles originating from the same subtomogram average structure of the prohibitin complex. An extra, matrix-facing density (yellow arrowhead) appears in Class 1. B. Subtomogram average structure of prohibitin particles belonging to Class 1 (gray transparent density) with fitted atomic models of prohibitin (PDB-9UNL, blue/orange) and the *m*-AAA protease, AFG3L2 (PDB-6NYY, pink). C. A triangulated surface mesh of the IMM depicted with subdomain classification and particles from class 1 (pink) and class 2 (blue) mapped back onto the surface. Gallery of cropped tomograms of prohibitin complexes from each class in two different subdomains (inner boundary membrane (IBM, yellow), and cristae body (CB, purple). Class 1 additional density denoted by yellow arrow. D. Voxel density scan of a prohibitin complex with additional matrix density. (top left) 2D slice of subtomogram average with line scan directionality indicated (Class 1 pink arrow, class 2 blue arrow). Additional peak from class 1 (pink line) noted by yellow arrow E. Quantification of particles from each class within each subdomain (left) and particle count normalized by total area of subdomain within each mitochondrion (right).

Mapping back the particles corresponding to the two distinct prohibitin structure classes showed that the majority of the particles from Class 1 are localized within the IBM, consistent with the overall enrichment of prohibitins found in this IMM subdomain (***Figure 2C*&*E***). This finding suggests that prohibitin-protease associations may occur preferentially within the IBM, where they could regulate protease activity on newly imported subunits of the OXPHOS machinery before they assemble in the cristae. This spatial coupling may provide an additional layer of quality control during respirasome formation and, consequently, mitochondrial metabolic activity. Interestingly, a subset of Class 1 particles localizes to the CBs and CJs, suggesting that prohibitin-mediated regulation of protease activity may also operate within these subdomains (***Figure 2E***). Defining how this context-specific structural variation is regulated and influences mitochondrial protein quality control will be an important area for future investigation.

### Prohibitin complexes are associated with unique membrane microenvironments

Prohibitins are essential for maintaining mitochondrial inner membrane architecture and lipid composition; however, how they influence — or are influenced by — their local membrane environment to support these functions remains poorly understood. To address this, we adapted a membrane patch-based approach we used previously to study the impact of co-translating ribosomes on the local membrane environment (Chang, Barad et al. 2025) to quantify local membrane ultrastructural parameters at prohibitin complexes. In brief, we identified the nearest triangle on the membrane surface mesh for each prohibitin complex and assigned all triangles within a radius of 15 nm around these triangles as the “prohibitin-associated” patch (***Figure 3A***). A radius of 150 Å was chosen based on measuring the radius of the subtomogram average structure of the prohibitin complex (∼100 Å) (***Figure 1C***) and extending it out 50 Å to ensure coverage of the full membrane ‘footprint’ occupied by the prohibitin complex (***Figures 3A***). Next, we used the *Surface Morphometrics* pipeline to measure phospholipid headgroup spacing, which we recently demonstrated can be a proxy for membrane thickness (Medina, Chang et al. 2025). We found that “prohibitin-associated” membrane patches exhibit a significant decrease in membrane thickness relative to a randomized patch control group across the IMM (***Figure 3B***), suggesting localized ‘thinning’ of the membrane at prohibitin complexes.

**Figure 3:**
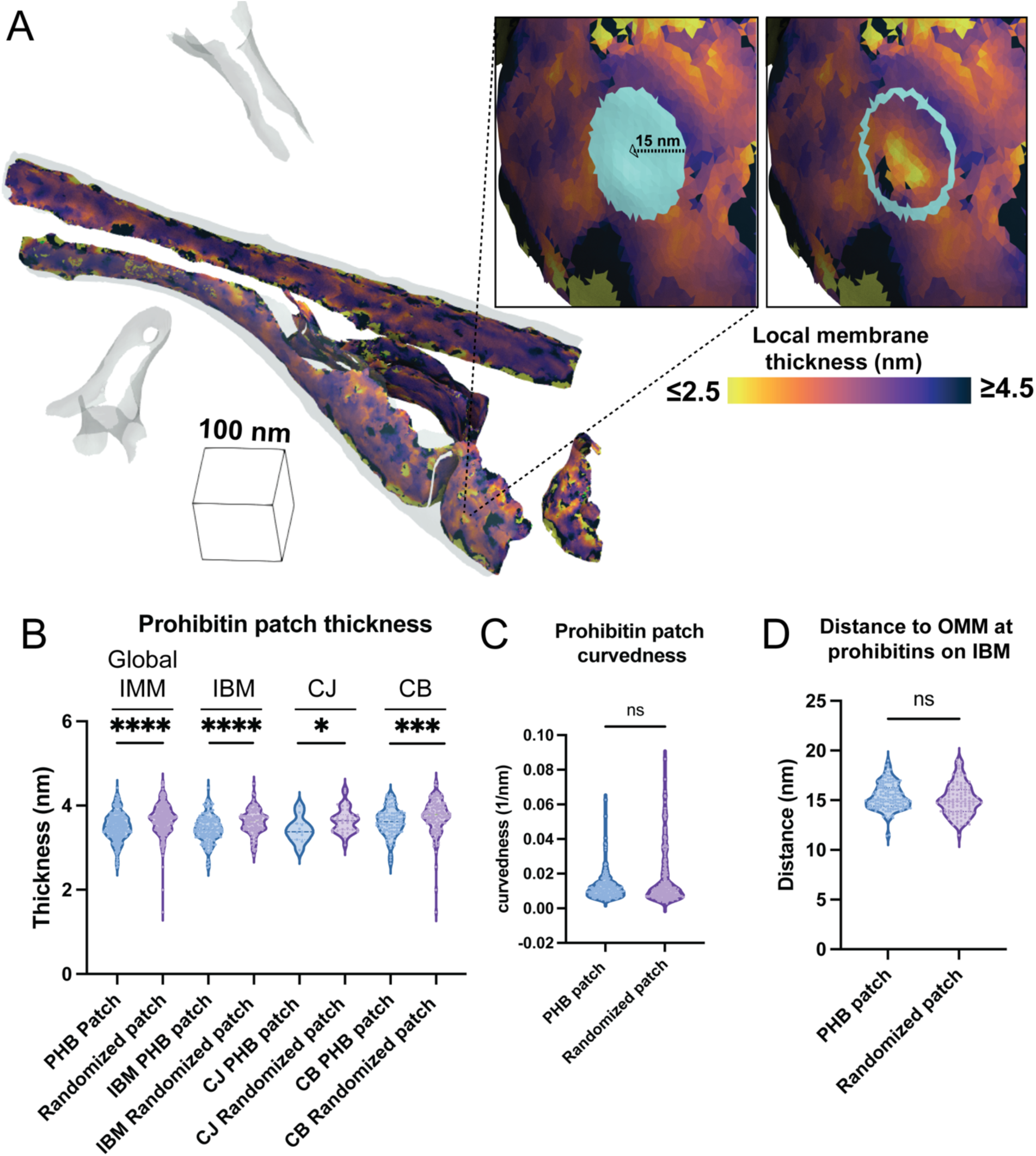
Prohibitin complexes induce local membrane remodeling. A. Per-triangle local thickness measurement overlaid on the IMM surface mesh reconstruction. Boxed insets show the region defined as the “prohibitin-associated” patch (PHB patch, teal) on the IMM surface mesh, as determined by assigning all triangles within a radius of 15 nm to the nearest triangle to the prohibitin complex on the membrane surface mesh. B. Quantification of patch thickness at prohibitin complexes (PHB patch) compared to a randomly generated patch (Randomized patch) along the global IMM and subdomains (IBM, CJ, CB). Quantifications from global IMM PHB patch n = 298, global IMM randomized patch n = 304, IBM PHB patch n = 186, IBM randomized patch n = 128, CJ PHB patch n = 11, CJ randomized patch n = 36, CB PHB patch n = 101, CB randomized patch n = 140. P values from Mann-Whitney U test are indicated *P< 0.05;***P<0.001, ****P<0.0001 C. Quantification of patch curvedness at prohibitin complexes (PHB patch) compared to a randomly generated patch (Randomized patch) along the IMM. Quantification from PHB patch n = 304, randomized patch n = 304. Ns = not significant D. Quantification of patch IMM to OMM distance at prohibitin complexes (PHB patch) compared to a randomly generated patch (Randomized patch) along the IMM. Quantification from PHB patch n = 144, randomized patch n = 100. Ns = not significant

We previously showed that different subdomains exhibit significant thickness variations across the IMM, with the IBM having the smallest lipid bilayer thickness (Medina, Chang et al. 2025). To control for these domain-specific differences that might explain the local thickness variations we observe, we separated our “prohibitin-associated” and randomized patches based on their IMM subdomain identity and compared membrane thickness between these groups. Strikingly, across all IMM subdomains, all prohibitin-associated patches showed significant decreases in membrane thickness compared with the random patch group. The nominal thickness values associated with “prohibitin-associated” patches vary slightly across domains (i.e., IBM = 3.42 nm; CJ = 3.49 nm; CB = 3.61 nm), yet in all cases, these patches are significantly thinner by approximately 0.2 nm compared to random patches in each corresponding subdomain. Notably, this difference is nearly twice as large as the differences in global membrane thickness across individual IMM subdomains (i.e., IBM = 3.65; CJ = 3.70 nm; CB = 3.8 nm) (Medina, Chang et al. 2025).

It has long been proposed that prohibitins may serve as a structural scaffold to recruit and stabilize unique lipid compositions, thereby forming functional microdomains within the IMM. Our finding of a significant reduction in membrane thickness locally at prohibitin complexes, regardless of the subdomain, provides direct evidence for localized differences in the lipid bilayer environment associated with prohibitin complexes. These results suggest two intriguing possibilities: (1) prohibitins actively remodel their local environment to induce membrane thinning, or (2) prohibitins preferentially localize to regions of local thickness minimum within the IMM. Future work is needed to determine whether these or other potential mechanisms exist to promote the observed local membrane changes associated with prohibitins.

Our finding that prohibitin complexes are associated with thinner membrane regions is also striking, given that we previously observed several other mitochondrial protein complexes, such as ATP synthase, are often associated with thicker membrane regions (Medina, Chang et al. 2025). We also had previously found a positive correlation between membrane thickness and curvature, a relationship absent in samples composed solely of in vitro-reconstituted phospholipid vesicles. This suggests that embedded proteins, particularly those containing extensive transmembrane helices, may contribute to this coupling. Consistent with this, recent reports from us and others show that other large inner membrane complexes, such as ATP synthase and respirasomes, are associated with local membrane domains of distinct curvature relative to the surrounding membrane environment under native cellular conditions. In the case of prohibitins, we observed no significant difference in local curvature between “prohibitin-associated” membrane patches and randomly selected patches (***Figure 3C***). We predict that differences in how these assemblies insert into the membrane likely underlie this distinction: whereas respirasomes contain multiple embedded transmembrane domains, each prohibitin subunit has only a single transmembrane helix positioned at the periphery of the ring. The central region of the lipid bilayer beneath the top of the prohibitin ‘dome’ contains few, if any, transmembrane helices, similar to *in vitro* reconstituted vesicles lacking embedded proteins, which may explain the lack of curvature in these regions.

Previous studies suggest that prohibitins, although localized to the inner membrane, may interact in *trans* with proteins on the outer mitochondrial membrane (Kasashima, Sumitani et al. 2008, Rose, Herrmann et al. 2025, Waltz, Righetto et al. 2025). Consistent with this, a recent study of prohibitins within intact *Chlamydomonas* cells revealed densities at the top of the prohibitin “dome” that appeared to contact the adjacent outer mitochondrial membrane (Waltz, Righetto et al. 2025). In contrast, we did not observe any bridging densities connecting the top of the prohibitin dome to the outer membrane in our subtomogram averages (***Figures 1 & 2***). This difference may reflect a limitation of subtomogram averaging, which converges on the most stable or abundant structural state and may therefore obscure transient or heterogeneous contacts. To probe whether such local interactions might occur across the entire population of prohibitin complexes in our dataset, we measured the distance between the outer and inner mitochondrial membrane at “prohibitin-associated” patches. Consistent with our reconstruction, we found no significant differences in the OMM-IMM spacing of “prohibitin patches” compared to random control regions (***Figure 3D***). These discrepancies may reflect differences in mitochondrial metabolic or functional state, or intrinsic species-specific differences between *Chlamydomonas* and mammalian cells.

## CONCLUSIONS AND PERSPECTIVES

Our study provides direct structural insights into the composition, localization, and local microenvironment of prohibitin complexes in the native context of MEF cells. We show that prohibitin complexes are IMS-facing, dome-like ring structures preferentially enriched in the IBM subdomain of the IMM. We show that a subset of prohibitin complexes associates with additional macromolecular complexes on the matrix-facing side of the inner membrane, which we propose corresponds to *m*-AAA proteases. This agrees with long-standing biochemical observations of prohibitin-protease interactions. Leveraging newly developed computational tools for quantitative membrane analysis, we further demonstrate that prohibitin complexes are associated with unique membrane environments characterized by reduced membrane thickness across all IMM subdomains. These findings provide direct experimental data for how the local membrane environment of prohibitins is organized to support their functions in lipid homeostasis and membrane integrity. The unique membrane and lipid environment associated with prohibitin complexes may serve as platforms to recruit proteins that bind either directly to prohibitin or directly to the lipid bilayer. Collectively, these studies provide a cellular context for elucidating the structural mechanisms underlying prohibitin function in mitochondrial biology. Given that dysregulation of prohibitins has been implicated in numerous human diseases (Theiss and Sitaraman 2011), including aging, neurodegeneration, cardiac disorders, and cancer, understanding these mechanisms has significant biomedical potential.

## ACKNOWLEDGEMENTS

We thank William Lessin at The Scripps Research Institute Hazen cryo-electron microscopy facility for microscope support. We thank Jean-Christophe Ducom and Lisa Dong at The Scripps Research Institute for computational support. We also thank R. Luke Wiseman, David DeRosier, and other members of the Grotjahn Lab for their critical input on the manuscript. M.M is supported by the Achievement Rewards for College Scientists (ARCS) Foundation fellowship. B.A.B. is supported by the Collins Medical Trust. D.A.G. is supported by The Pew Scholars Program, The Bachrach Family Foundation, Nadia’s Gift Foundation Innovator Award of the Damon Runyon Cancer Foundation (DRR-65-21), and the National Institutes of Health (NIH) grant RF1NS125674. This work used equipment supported by NIH grant S10OD032467. Assistance from ChatGPT-4.0 (Open AI, https://chat.openai.com/) and Grammarly’s AI tool was utilized to improve the clarity, grammar, and conciseness of the manuscript text.

## AUTHOR CONTRIBUTIONS

**M. Medina:** Conceptualization, Data curation, Formal analysis, Investigation, Methodology, Software, Validation, Visualization, Writing - original draft, Writing - reviewing and editing.

**H. Rahmani:** Data curation, Formal Analysis, Investigation, Methodology, Software, Validation, Writing - reviewing and editing.

**Y.-T. Chang:** Data curation, Formal analysis, Methodology, Software, Validation, Visualization, Writing - reviewing and editing.

**B. A. Barad:** Conceptualization, Methodology, Software, Writing - reviewing and editing.

**D. A. Grotjahn:** Conceptualization, Funding acquisition, Methodology, Project administration, Resources, Supervision, Visualization, Writing - original draft, Writing - reviewing and editing.

## METHODS

### Sample Preparation and cryo-ET data collection

Sample preparation and cryo-ET data collection were performed as previously described (Medina, Chang et al. 2025). In brief, mouse embryonic fibroblasts with mitochondria labeled GFP (MEFmtGFP) (Wang, Wang et al. 2012) were cultured on UV sterilized R ¼ Carbon 200-mesh gold electron microscopy (EM) grids (Quantifoil Micro Tools) and vitrified using a Vitrobot Mark 4 (Thermo Fisher Scientific). Cryo-focused ion beam (cryo-FIB) milling of lamellae was performed using an Aquilos dual-beam cryo-FIB/SEM instrument (Thermo Fisher Scientific). Targeted cells were milled using an automated cryo-FIB milling workflow (Buckley, Gervinskas et al. 2020) and xT software with MAPS (Thermo Fisher).

EM grids containing lamellae were transferred into a 300keV Titan Krios microscope (Thermo Fisher Scientific), equipped with a K3 Summit direct electron detector camera (Gatan), and a BioQuantum energy filter (Gatan). Tilt series were acquired using parallel cryo-electron tomography (PACE-tomo) (Eisenstein, Yanagisawa et al. 2023) in SerialEM (Mastronarde 2003). Tilt series were acquired at magnification 53,000x with a pixel size of 1.662 Å and a nominal defocus range between (−4 to −8 µm). Data collection was done in a dose symmetric scheme with 2° steps between −60° and +60° centered on −11° pretilt angle. The total dose per tilt was 3.00 e/Å2, and the total accumulated dose for the tilt series was under 123 e/Å2.

### Tilt series processing and reconstruction

Tilt series processing and reconstruction were performed as previously described (Medina, Chang et al. 2025). Dose fractionated tilt series micrograph movies were subjected to CTF estimation and motion correction in Warp (Tegunov and Cramer 2019). Tilt series alignment was performed in Etomo using post-polishing platinum sputter beads as fiducials. The reconstruction was done using Warp (Tegunov and Cramer 2019) back projection to produce 4, 6, and 8 times binned tomograms, corresponding to pixel sizes of 6.65Å, 9.98Å, and 13.3Å, respectively. These data have been previously deposited in the Electron Microscopy Public Image Archive under accession code EMPIAR-13056.

### Prohibitin particle picking and subtomogram averaging

Prohibitin complexes were picked using a two-point directional picking strategy in the software package i3 (Winkler, Zhu et al. 2009) on tomograms binned to 4 (6.65 Å), which were post-processed with a median filter in the z direction with a kernel size of 11 pixels and an MTF filter (***Supplementary Figure 1***). Each particle was denoted by two points: the first one on the IMM and the second one on top of the prohibitin complex. These points were used to define a vector in 3D, and in-house Python scripts were used to align all vectors to the z-axis and apply a random rotation around it. This rotation helps mitigate misalignment due to the missing-wedge artifact when particles are aligned to the reference. These particles were then used as input for alignment and classification in i3 package and then further refined in RELION (Zivanov, Otón et al. 2022). To focus on lower-resolution information and expand the box size, we used tomograms reconstructed at a pixel size of 9.98 Å for subtomogram averaging. The particles were first aligned translationally to the global average and then classified into four classes in i3. After initial alignment, the particles were classified using a cylindrical mask that covered the area under the prohibitin complex within the mitochondrial matrix, yielding two structural classes (Class 1 and Class 2). The i3 alignment files were then converted to RELION star files using in-house Python scripts, and the subvolumes were extracted at a pixel size = 3.32 Å in Warp. All particles and those corresponding to Class 1 and Class 2 were refined separately in RELION-4.0 using relion-autorefine. ArtiaX module in Chimerax was used to visualize these particles in the original tomograms (Ermel, Arghittu et al. 2022).

### Membrane surface generation and morphometrics analysis

Membrane surface generation and analyses using the *Surface Morphometrics* pipeline were performed as previously described (Medina, Chang et al. 2025). In brief, membranes were segmented using Membrain-Seg (Lamm, Zufferey et al. 2024) before manual labeling of distinct membranes (i.e., IMM, OMM, etc.) in AMIRA (Thermo Fisher Scientific). The voxel segmentation membrane label files were then exported from AMIRA and input into Surface Morphometrics for surface generation. Surfaces were subdivided into individual segments based on the connected components of the membrane graph to get “per-component” analyses to establish reasonable estimates of independent samples within each tomogram. Each membrane surface was processed using *Surface Morphometrics* to obtain measurements of membrane distance, curvature, orientation, and thickness.

### Prohibitin IMM subdomain assignment

The IMM subdomains for each IMM surface were classified based on the OMM-IMM distances, as previously described (Medina, Chang et al. 2025). The RELION star file containing the particle coordinates, alignment, and classification information from i3 was used to determine the nearest triangle on the IMM surface and assign a ‘subdomain identity’ to each prohibitin particle. The ‘subdomain identity’ and total area of the corresponding IMM subdomain were added to the RELION star file as new columns. The resulting RELION star file was used to calculate the number of prohibitins in each subdomain with and without normalization. Each prohibitin particle was visually inspected to verify correct subdomain assignment using ArtiaX (Ermel, Arghittu et al. 2022).

### Prohibitin patch designation and morphometric analyses

PHB coordinates were obtained from the RELION star files corresponding to each tomogram in a similar manner used in (Chang, Barad et al. 2025). The IMM surface coordinates and thickness for each mitochondrion were from triangle graph files (.gt) generated by *Surface Morphometrics* pipeline. The nearest IMM triangle in the IMM surface mesh reconstruction to each PHB particle coordinate was identified using a k-dimensional tree Python function. To avoid cross-assignment of particles from other surfaces in the same tomogram, we excluded any nearest IMM triangles located farther than the height of a PHB particle (9 nm). The remaining nearest IMM surface triangles were designated as “patch centers.” Around each patch center, we searched for triangles within a 15 nm radius to define the PHB-associated patch. Randomized PHB-associated patches for each mitochondrion were generated based on the following criteria: (1) the number of randomized patches matched the number of PHB-associated patches, and (2) the distances between randomized patch centers were greater than 15 nm. This process was performed using “find_IMM_patches_for_PHB.py”. The average thickness for each PHB-associated patch and each randomized patch were calculated by “average_thickness_calculation_per_patch.py” and visualized as violin plot. The Mann–Whitney U test was applied to assess the statistical significance of differences. The generation of the violin plot and statistic test were performed using Prism (GraphPad).

PHB-associated patches from multiple mitochondria were extracted as individual patch surfaces, preserving the coordinates and normal vectors of each surface triangle, using the script “extract_single_patch.py”. Before performing line scanning, the normal vectors of the surface triangles in each patch were curated to ensure they pointed in the same direction as the vector from the patch center to the corresponding PHB particle center. These curated patch surfaces were then correlated with the tomogram and served as the reference points for the line scanning process. For each PHB, a line scanning density profile was generated by sampling tomogram intensity values along the direction of the curated normal vectors. The scan extended from −20 nm (toward the matrix) to +20 nm (toward the IMS and OMM) with 0.25 nm steps. The average PHB line-scanning density profile was obtained by averaging the intensity profiles across all prohibitin particles after normalizing each scan to the maximum density value of the IMM.

## Data and Code availability

All tilt series, reconstructed tomograms, voxel segmentations, and reconstructed mesh surfaces used for quantifications were previously deposited in the Electron Microscopy Public Image Archive (EMPIAR) under accession code EMPIAR-13056. All subtomogram averages were deposited in the Electron Microscopy Data Bank (EMDB) under accession codes EMD-XXXX. All scripts used for Surface Morphometrics are available at https://github.com/grotjahnlab/surface_morphometrics).

