## Supplemental Figure for "Prohibitin complexes associate with unique membrane microdomains in cells"

**SUPPLEMENTARY FIGURE**

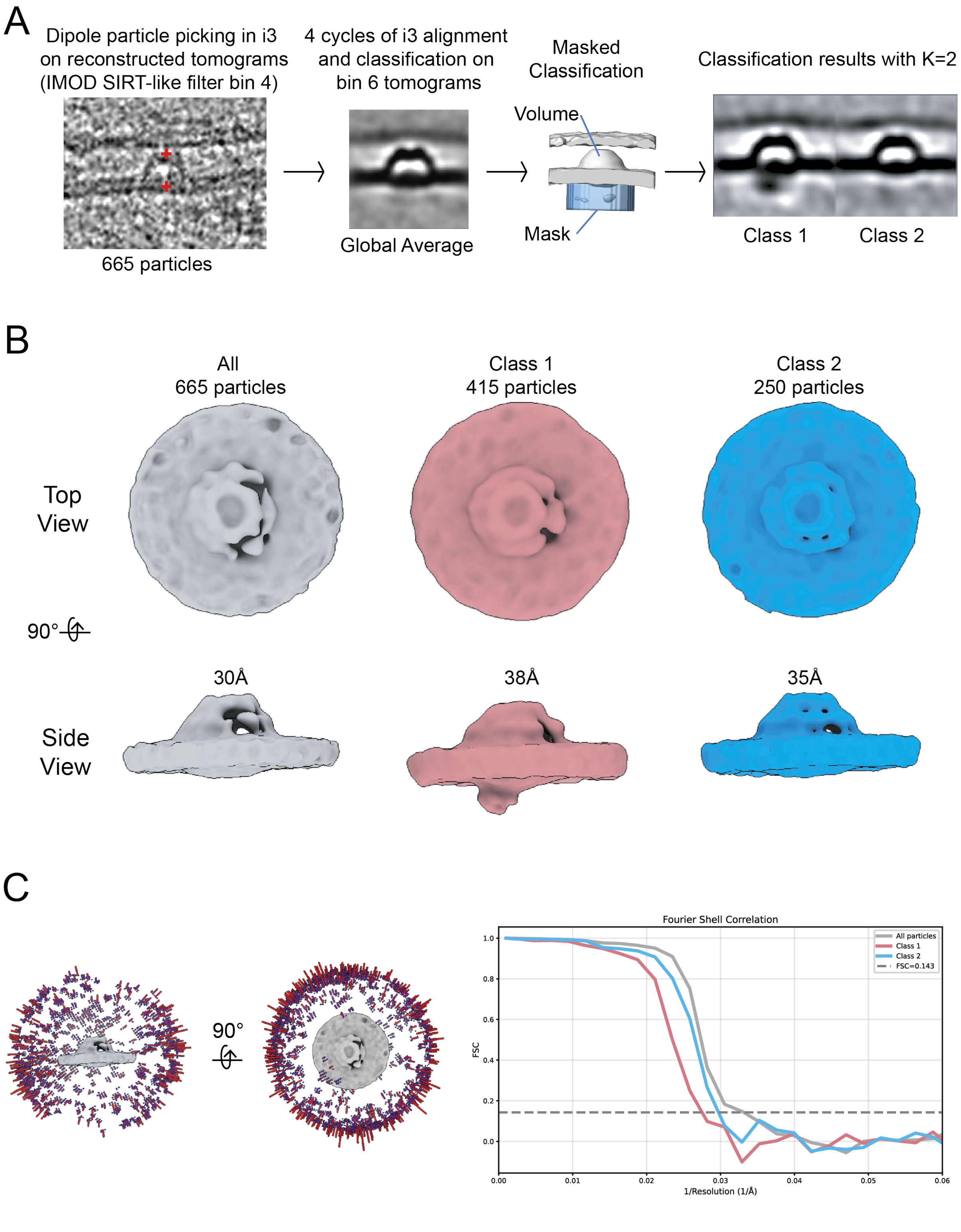

**Supplemental Figure 1. Subtomogram averaging workflow**

A. Workflow for sub-tomogram averaging for prohibitin complex from all 665 particles in (gray) and subclassification.

B. Subtomogram average structure from all prohibitin particles (Grey, 665 particles) and the subclasses (Class 1 – pink, Class 2 – blue).

C. (left) The final Euler distribution of the prohibitin structure from all 665 particles from RELION refinement. (right) FSC curves from all 3 refinements in (B) from RELION.
